# Can deep learning provide a generalizable model for dynamic sound encoding in auditory cortex?

**DOI:** 10.1101/2022.06.10.495698

**Authors:** Jacob R. Pennington, Stephen V. David

**Author notes:** Correspondence: SVD, 3181 SW Sam Jackson Park Road, MC L335A, Portland, OR 97239.

## Abstract

Convolutional neural networks (CNNs) can provide powerful and flexible models of neural sensory processing. However, the utility of CNNs in studying the auditory system has been limited by their requirement for large datasets and the complex response properties of single auditory neurons. To address these limitations, we developed a population encoding model: a CNN that simultaneously predicts activity of several hundred neurons recorded during presentation of a large set of natural sounds. This approach defines a shared spectro-temporal space and pools statistical power across neurons. Population models of varying architecture performed consistently better than traditional linear-nonlinear models on data from primary and non-primary auditory cortex. Moreover, population models were highly generalizable. The output layer of a model pre-trained on one population of neurons could be fit to novel single units, achieving performance equivalent to that of neurons in the original fit data. This ability to generalize suggests that population encoding models capture a general set of computations performed by auditory cortex.

## Introduction

A complete understanding of neural sensory processing requires computational models that can account for brain activity evoked by arbitrary natural stimuli^1^. In the auditory cortex, encoding models such as the widely used linear-nonlinear spectro-temporal receptive field (LN model) can account for sound-evoked spiking activity in some neurons, but often fail to predict time-varying responses to complex stimuli such as natural sounds^2,3^. Encoding models are used to study auditory coding in many neurophysiological signals beyond single neuron spikes, including calcium imaging^4^, spiking ensembles^5^, human LFP^6,7^, MEG^8^, and fMRI BOLD^9,10^. Thus, improved encoding models are of broad value to research on the auditory system. However, there is currently no consensus on what model architecture should replace the LN model as the standard for auditory neurons.

Variants of the LN model have been proposed that provide a more accurate characterization of auditory coding. Some of these variants build on traditional systems identification methods, accounting for second- and higher order nonlinearities^11,12^. Others have incorporated nonlinear elements derived directly from biological circuits, like short-term synaptic plasticity and gain control from local inhibitory populations^3,13,14^. A third approach has been to use artificial neural networks, generalized models that combine large numbers of linear-nonlinear units^15^. While theoretically appealing, large neural networks have proven challenging to fit, especially when data set size is limited. Direct comparisons between these different methods remain limited, as new models are typically compared only with the LN model as a baseline (but see^6,14,15^).

In the current study, we explored convolutional neural networks (CNNs) as a method for improving upon existing encoding models of neural spiking data. Advances in machine learning, in particular the development of backpropagation algorithms for CNNs, have opened up the possibility of applying neural network analysis to electrophysiology datasets^6,16^. CNNs have been adopted widely for signal processing problems, including speech recognition and other acoustic analysis^17,18^. A small number of studies have indicated that CNNs can describe human auditory BOLD fMRI data^16^ and ECoG data^6^. In addition, CNNs have been applied to models of natural image representation in visual cortex^19^. It remains an open question whether CNNs can provide a useful characterization of single-neuron activity in auditory cortex.

One particular appeal of CNNs is that they can serve as “foundation models,” pre-trained on one task but then applied to a wide range of new problems^20^. Such approaches are widely used for machine learning problems, including auditory^21^ and visual signal processing^22^. Pre-trained CNNs have also been useful for analyzing neural data when limited data set size prevents fitting large models directly^23^. Motivated by the success of this approach, we argue that an effective CNN-based encoding model should be fully generalizable. That is, a CNN that completely describes neural sensory processing should account for the encoding properties of neurons that were not included in the original model fit. Moreover, such a model should predict higher-order properties of neural activity, like sparse coding, that are not captured by traditional LN models but are characteristic of cortical sensory activity^24,25^.

In order to fit CNN models and evaluate their generalizability, we recorded the time-varying spiking activity of a population of single neurons in auditory cortex during presentation of a large natural sound library. To leverage statistical power across neurons, we developed a population encoding model, in which the activity of many neurons is predicted by a single CNN with input layers shared across neurons. We found that these population models accounted for cortical activity substantially better than traditional LN models. In addition, pre-trained population models successfully generalized to novel neural data and more closely reproduced the sparse coding properties of auditory cortex. Thus, CNN models fit with large neural populations and diverse stimuli can provide a generalizable model of sound encoding by auditory cortex.

## Results

To characterize the neural encoding of natural sounds, we recorded spiking activity from auditory cortex of awake, passively listening ferrets during presentation of a diverse set of natural sound samples. Neural data was collected with 64-or 128 channel linear silicon arrays that recorded simultaneous activity of 10-95 neurons across multiple cortical laminae during each experiment. Recordings were performed in primary auditory cortex (A1) and a secondary auditory field (PEG) located anterior-ventral to A1^26,27^. The same stimuli were presented in all recordings (A1: 22 recording sites, 849 units; PEG: 11 sites, 398 units; 5 animals).

### Convolutional neural networks with a shared tuning space for neural populations

We used a convolutional neural network (CNN) to describe the functional relationship between the natural sound spectrogram and the time-varying spiking activity (Fig. 1). Machine learning tools have proven effective for studying a wide range of analytically similar problems. However, fitting these models requires large datasets, and the amount of data available from many neurophysiological studies is limited. This limitation is compounded by the fact that sensory-evoked neural activity is not reliable, varying substantially across repeated stimulus presentations. The current study took advantage of the fact that identical stimuli were presented during multiple experiments to fit CNNs that simultaneously modeled the entire population of neurons being studied. In this framework, the stimulus spectrogram provides input to a series of fully connected layers that are shared across all neurons. The final layer computes separate weighted sums to predict the activity of each neuron individually (Fig. 1D-F). Thus, the earlier layers comprise a shared, general model of sound processing in auditory cortex which is mapped to individual responses only in the final layer.

**Figure 1.**
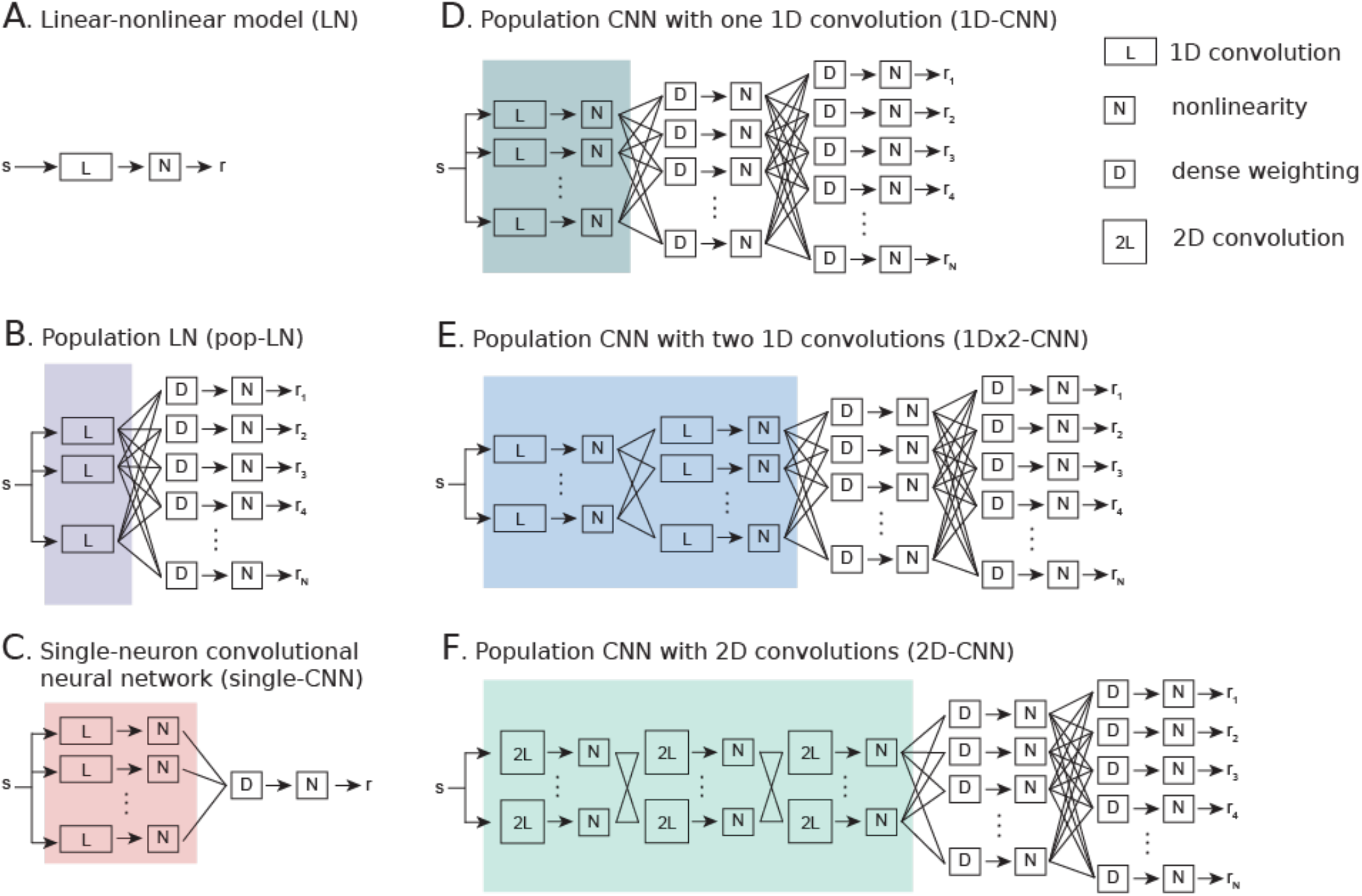
Convolutional neural networks (CNNs) provide a natural extension of standard linear-nonlinear (LN) models of auditory encoding. **A**. LN model consists of a single convolutional filter (L) followed by a static nonlinearity (N). This convolution is one-dimensional: a separate filter is convolved in time for each frequency channel, then the results are summed across frequency. Arrows indicate passthrough of intermediate computations between units within a layer. **B**. Population LN (pop-LN) model is composed of a bank of convolutional units (purple shading) followed by one dense unit (D) and static nonlinearity per neuron. Lines represent weightings between layers. **C**. Single-neuron convolutional neural network (single-CNN) consists of a bank of LN units (red shading) linearly combined in a subsequent dense unit and followed by another static nonlinearity. **D-F**. Population CNN models consist of one or more convolutional layers and two fully connected layers. Dense units in the final layer generate the output for each neuron. Convolutional units are either one-dimensional (1D-CNN) or two-dimensional (2D-CNN). The 2D-CNN model includes three convolutional layers in sequence (**F**, light green shading), while the 1D-CNN model either has one convolutional layer (**D**, dark green shading) or two (1Dx2-CNN, **E**, blue shading).

Within the population model framework, we compared three CNN architectures (Fig. 1D-F) to the standard LN architecture (Fig. 1A), which is widely used to describe the encoding properties of single neurons^14,28,29^. CNNs used in visual processing problems typically apply multiple layers of small, two-dimensional (2D) convolutional kernels to an input image^19^. We adapted this architecture to an auditory model, in which 2D filters are convolved along the time and frequency dimensions of the sound spectrogram (2D-CNN, Fig. 1F)^6^. To compare the CNN approach directly to LN models typically used in the auditory system, we also developed a one-dimensional CNN in which filters are convolved only in time. For these 1D models, each filter is convolved in time with a weighted sum of either the input stimulus channels or the outputs of a previous layer. We evaluated two versions of this architecture: one with a single convolutional layer (1D-CNN, Fig. 1D) and one with two convolutional layers in sequence (1Dx2-CNN, Fig. 1E).

We also implemented two intermediate architectures to control for differences between population CNN models and the standard LN model. To control for increased statistical power gained by pooling data across neurons in population models, we fit a population LN model (pop-LN, Fig. 1B), in which the simultaneous activity of the recorded neural population is modeled as the weighted sum of a shared bank of 1D convolutional filters followed by nonlinear rectification. To distinguish possible benefits of the population model approach from benefits of the neural network architecture, we fit separate CNN models for each neuron in the dataset (single-CNN, Fig. 1C). To accommodate sampling limitations of single unit data, these models contained substantially fewer units than the larger population models.

All models were fit using standard back-propagation methods^30^, which minimized the mean-squared error between predicted and actual time-varying neural activity. Fitting was carried out in two stages. First, parameters for the entire model were fit for all neurons simultaneously. Second, weights in the final layer were re-fit for each neuron individually (see Methods). Fitted models were then used to predict the response evoked by stimuli in a validation dataset that was not used for fitting (Supplementary Fig. 1). Model performance was evaluated on this separate dataset by measuring prediction correlation, the noise-corrected Pearson correlation coefficient between predicted and actual time-varying response, for each neuron^31,32^. A value of 1.0 indicates that a neuron’s activity was predicted as accurately as possible given the uncertainty in the actual response, and a value of 0 indicates chance performance.

An example of a 1Dx2-CNN fit to the A1 dataset illustrates the comprehensive nature of the population CNN models (Fig. 2). The first layer consists of 70 1D convolutional units that resemble the filters used in standard LN model fits (Fig. 2D, E). This layer, in conjunction with the 80-unit convolutional layer that follows, defines a space of spectro-temporal channels upon which the subsequent dense layers depend. The positive and negative weights connecting these dense layers produce the time-varying output specific to each unit (Fig. 2C), which is compared to the actual neural activity (Fig. 2B). Prediction correlation varied across the neural population but was close to a maximum value of 1.0 for many units (Fig. 2F).

**Figure 2.**
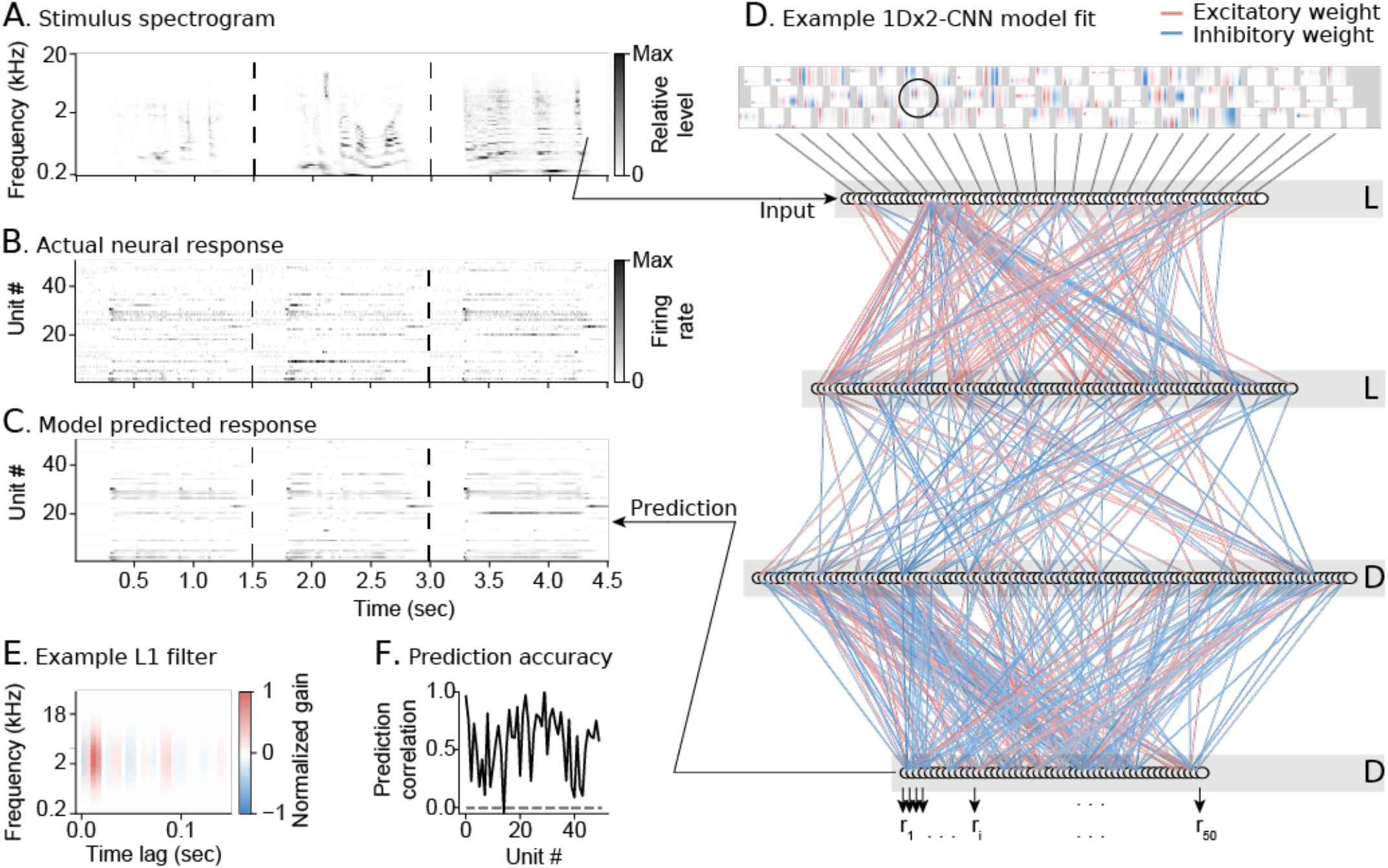
Example population CNN model fit for 849 A1 neurons. **A**. Example spectrogram of three natural sounds in the validation set used to measure model performance. **B**. Heat map shows the actual time-varying spike rate of one neuron per row, for n = 50 / 849 neurons from the A1 dataset. Spike rate is normalized for each neuron between 0 (white) and maximum (black). **C**. Heat map of predicted activity for the same units, plotted as in B. **D**. Schematic of model layers with fitted weights. 1D filters in the first two layers (top) establish a shared set of spectro-temporal channels. Line color connecting subsequent layers indicates the weight, with positive (excitatory) weights in red and negative (inhibitory) weights in blue. Each unit in the output layer (bottom) predicts the activity of one neuron **E**. Example filter from the first layer of the population CNN model (circled in D). The filter resembles one typically observed in a standard LN model for a single neuron. **F**. Prediction correlation for each unit shown in B-C.

### CNN models consistently outperform LN models

For each model architecture, we fit variants of differing complexity by changing the number of units in one or more layers, which in turn varied the number of fit parameters. Note that a single-CNN model with one filter in the first layer reduces to a standard LN model. Thus, by varying layer size, we explored a continuous space of models ranging from the LN model to much larger CNNs. For the LN model, we varied complexity by changing the rank of the convolutional filter^31^.

First, we consider the results from A1 data. We compared the performance of models with different architectures and sizes in a Pareto plot (Fig. 3A). Prediction correlation increased as model size increased and approached asymptotic performance at around 150-200 free parameters per neuron for most architectures. We selected an exemplar model from each architecture with performance and complexity in this range (circled points, Fig. 3A). Unless otherwise noted, subsequent analysis focuses on these models and a subset of auditory-responsive neurons, defined as neurons for which activity was predicted above chance by the 1Dx2-CNN, pop-LN and single-CNN exemplar models (p < 0.05; A1: n = 777/849, PEG: n = 339/398).

**Figure 3.**
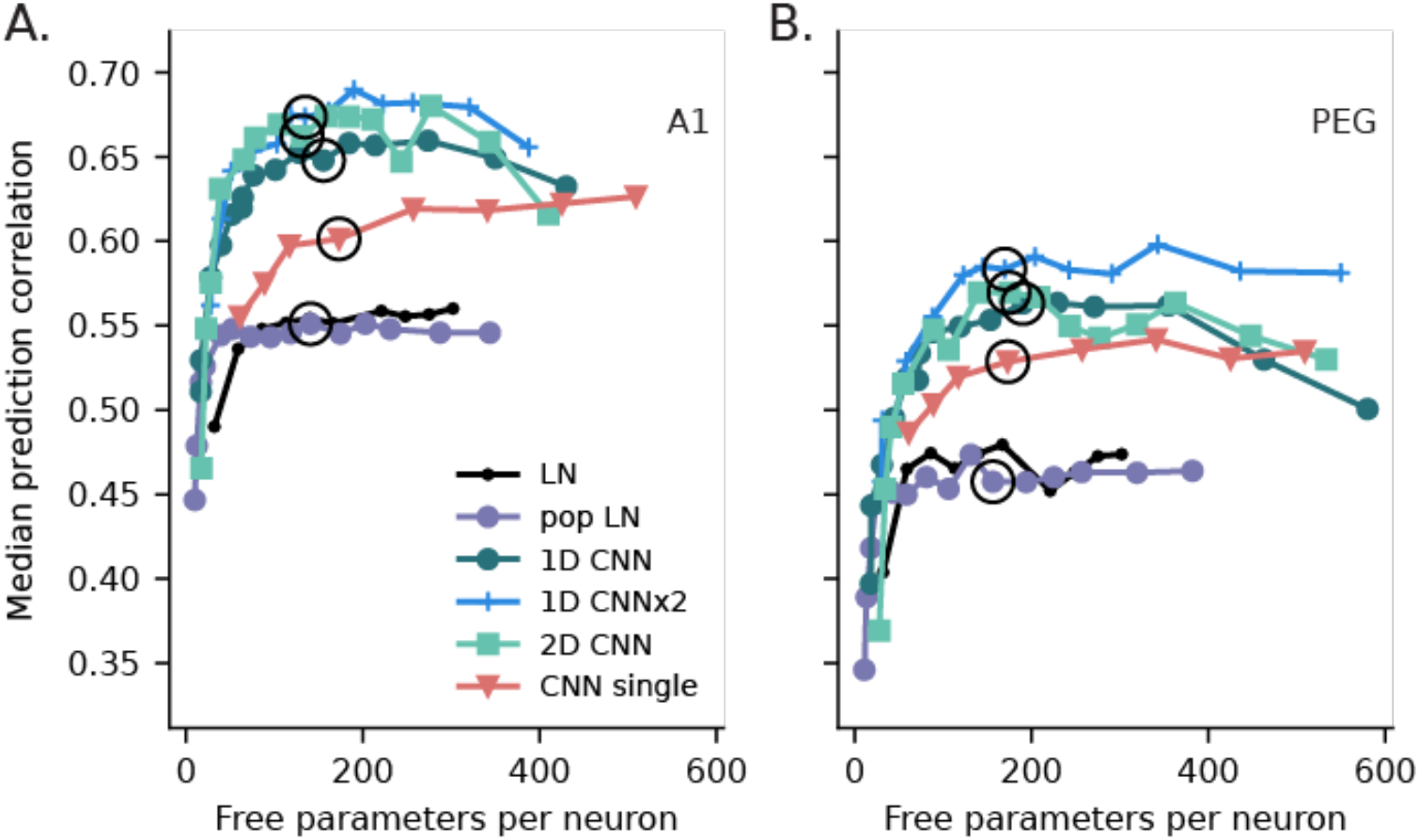
Performance of models from each architecture with variable parameter count. **A**. Pareto plot compares model complexity (parameter count) versus median prediction correlation for each model in A1 (n = 777 auditory-responsive neurons). CNN models maintained consistently higher prediction correlation across a wide range of complexity. Lines connect models in each of the six architectures. Circles indicate exemplar models from each category with similar complexity, which are examined in more detail in subsequent analyses. For population models, parameter count is normalized by the number of neurons that were simultaneously fit. **B**. Pareto plot comparing model performance in a secondary field (PEG, n = 339), plotted as in A. Relative performance differences across model types were comparable to A1, but median prediction correlation was lower for all models.

When model complexity was matched (i.e., for exemplar models with the same number of fit parameters per neuron), prediction correlation was higher for all population CNN models (1D-CNN, 1Dx2-CNN, 2D-CNN) than for the other architectures (Figs. 3-4, Supplementary Fig. 1). The greater accuracy of CNN models was consistent across the neural population: the 1Dx2-CNN model predicted responses more accurately than the pop-LN model (p < 0.05, jackknifed t-test) for almost half of auditory-responsive A1 neurons (376/777) while the opposite was true for only four neurons (Fig. 4A). On average, the best 1Dx2-CNN model accounted for 47% of the explainable variance in the A1 data, compared to 31% for the best LN model and 39% for the best single-CNN model. The explanatory power of the 1Dx2-CNN model was also greater than that of a nonlinear model that explicitly incorporates short-term plasticity and gain control, which accounted for 37% of explainable variance in a similar natural sound dataset^14^.

**Figure 4.**
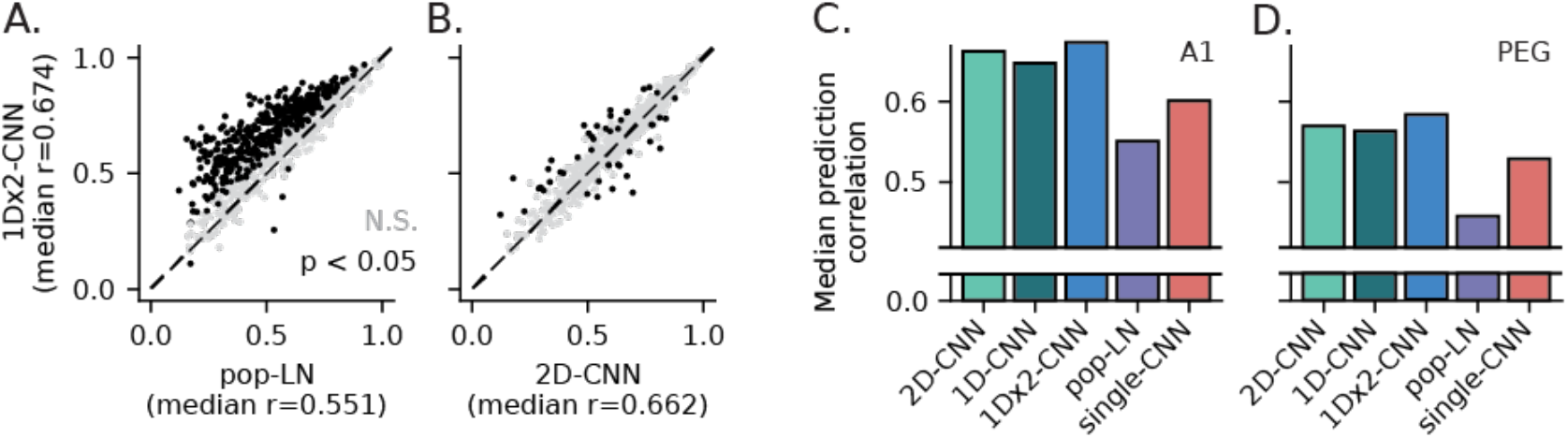
LN and CNN exemplar model performance. **A**. Scatter plot comparing prediction correlation for exemplar pop-LN and 1Dx2-CNN models for each neuron in the A1 dataset. The 1Dx2-CNN model had significantly higher prediction correlation for 376 of the 777 neurons. **B**. Scatter plot comparing the 2D-CNN and 1Dx2-CNN models, plotted as in A. Prediction correlations were comparable in this case, but the 1Dx2-CNN model still represents a small overall improvement (signed-rank test, p = 1.10 × 10^−8^). **C**. Median prediction correlation in A1 for exemplar models representing each architecture. All differences were statistically significant (signed-rank test, 1D-CNN vs. 2D-CNN: p = 9.21 × 10^−3^, other comparisons: p < 10^−7^). **D**. Prediction correlation for PEG data, plotted as in C. For this dataset, the difference between 1Dx2-CNN and 2D-CNN was not significant (signed-rank test, p = 0.883), but all other differences were (p < 10^−4^). Although overall prediction correlation was lower for PEG than A1, the relative difference in performance between models was the same for both areas, and the 1Dx2-CNN model was the best-performing in both.

In addition to their increased accuracy relative to LN models, the population CNN models performed better than the single-CNN model. This result indicates that the improved performance of the population models was not solely due to their neural network architecture but also reflected the gain in statistical power from pooling data across neurons. At the same time, the single-CNN model did outperform both the LN and pop-LN models, confirming a benefit of the CNN architecture over the traditional LN model. The single-CNN architecture also continued to increase in accuracy as parameter count grew, suggesting that a large CNN can indeed be an effective single-neuron model if a sufficiently large dataset is available.

Among population CNN models of similar complexity, prediction correlation was quite similar, suggesting that the specific architecture was not critical to performance (Fig. 4B, C). The detailed dynamics predicted by each CNN model were also closely matched (Supplementary Figs. 1, 2). However, adding an extra convolutional layer to the 1D-CNN model (Fig. 1D) did increase prediction correlation (signed-rank test, p = 3.89 × 10^−18^, Fig 4C), making 1Dx2-CNN (Fig. 1E) the best-performing model tested. This modest but significant difference suggests that further improvements can be achieved by testing additional CNN architectures.

Median prediction correlation for the LN and pop-LN models was nearly identical, indicating that pooling data across the neural population did not benefit performance of this less complex architecture (signed-rank test between exemplar models; A1: p = 0.133, PEG: p = 0.167). We confirmed that the two models also produced very similar predictions (Supplementary Fig. 2). For subsequent analyses, we focused on the pop-LN model, since there are fewer differences between its architecture and the population CNNs.

The relationships between models described for A1 also held true for the PEG data (Figs. 3B, 4D). However, median prediction correlation was consistently lower for PEG across all architectures and model sizes. This difference persisted after controlling for reduced response reliability (SNR) of PEG responses relative to A1 (Supplementary Fig. 3).

### Population models generalize to novel datasets

We hypothesized that, when fit to the activity of many neurons, population models capture an encoding subspace shared across neurons in the brain area that they describe. Whether constrained by information bottlenecks in neural circuits or by developmental plasticity following exposure to behaviorally important sound features, the space of sound representations in cortex is likely to be lower-dimensional than the space of all possible stimuli. Similarly, the shared tuning described by population models is constrained by the dimensionality of the final hidden network layer. We reasoned that if these models captured the actual neural subspace, then they should be able generalize to data from neurons that were not included in the original model fit.

To test this hypothesis, we re-fit the 1Dx2-CNN, pop-LN, and single-CNN models using two alternative approaches (Fig. 5A). In a “held-out” model, all data from one recording site was excluded during the first stage of the model fit. The output layer was then re-fit to each excluded neuron in the second stage, but model parameters were kept fixed for all preceding layers. In other words, the held-out model was pre-trained on data that excluded both the neuron being predicted and its neighbors from the same recording site. Excluding the entire recording site precluded the possibility of fitting to activity of neurons in the local network with highly overlapping selectivity. As a control, we also fit a “matched” model in which a subset of neurons from other recording sites was excluded during the first stage of fitting, such that the number of neurons excluded was the same as for the held-out model. This design ensured that the amount of fit data was matched to that of the held-out model, and data from the predicted neuron was used to fit all model layers (see Methods).

**Figure 5.**
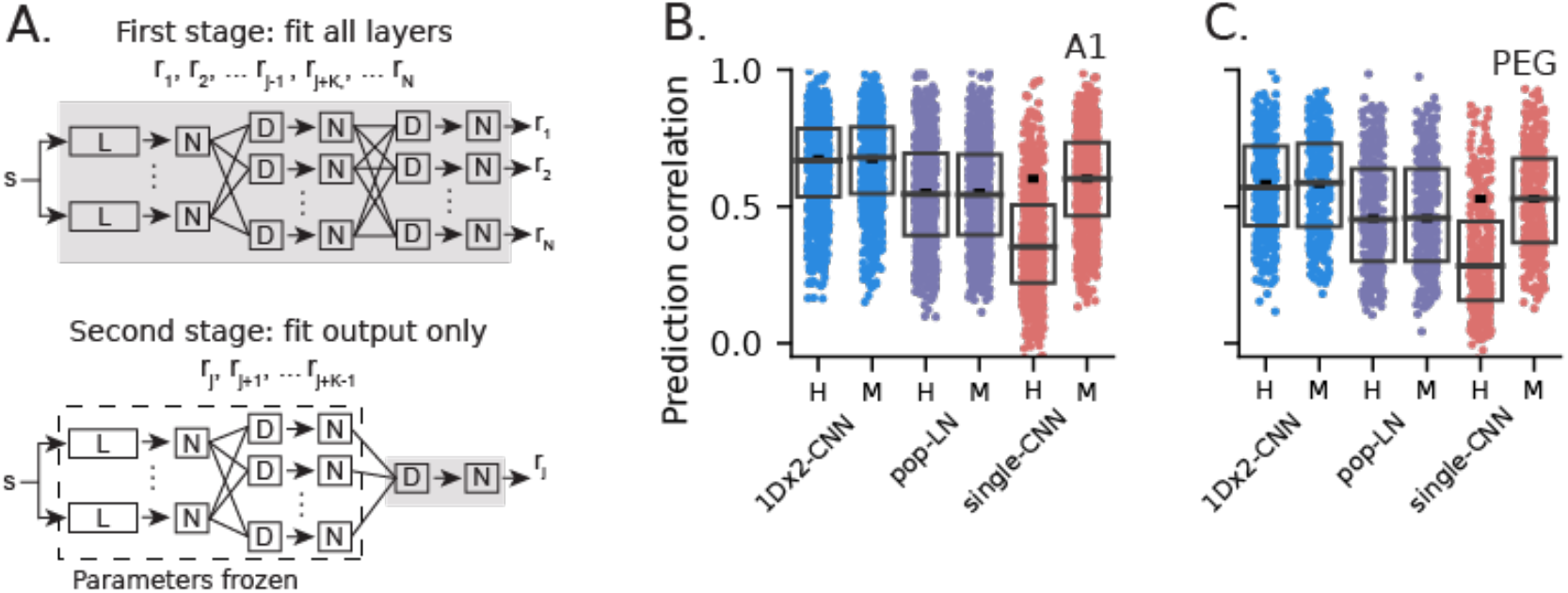
Generalization of population models to novel neural data. **A**. Schematic of two-stage fitting process for held-out and matched models. For the held-out model, responses of all K neurons from a recording site were excluded from the first stage fit. For the matched model, K neurons from other sites with similar prediction correlation were excluded during the first stage. For both models, all parameters except those in the final layer were frozen (dashed blue box) while individually fitting the K excluded responses in the second stage. **B**. Prediction correlation of A1 held-out (H) and matched (M) models. Boxes show the 1st, 2nd and 3rd data quartiles, and the small horizontal dash shows median performance for the full model (Fig. 4C). For both population models, the difference in prediction correlation between held-out and matched was not significant (signed-rank test, 1Dx2-CNN: p = 0.751; pop-LN: p = 0.866), indicating that these models generalize well to novel data. In contrast, there was a substantial decrease in prediction correlation for the held-out single-CNN model (p = 4.93 × 10^−81^). **C**. Generalization for PEG neurons, plotted as in B. Once again, performance was the same for held-out and matched population models but significantly decreased for the held-out single-CNN model (signed-rank test, 1Dx2-CNN: p = 0.418; pop-LN: p = 0.941; single-CNN: p = 6.46 × 10^−33^).

If performance of the held-out and matched versions of a model is equal for a given neuron, then the held-out model already accounts for the spectro-temporal encoding properties of that neuron in its pre-trained layers. In this case, we can say that the model generalizes well: the response of any novel neuron from the recorded brain area can be accounted for simply by re-fitting the output layer of a pre-trained model. When we compared performance of the held-out and matched versions of the three models (Fig. 5B, C), we found that the 1Dx2-CNN and pop-LN models both generalized well: there was no difference in prediction correlation between the two fitting strategies (signed-rank test, A1; 1Dx2-CNN: p = 0.751; pop-LN: p = 0.866; PEG; 1Dx2-CNN: p = 0.418; pop-LN: p = 0.941). The generalizability of the 1Dx2-CNN model also extended across brain regions (Supplementary Fig. 3). In contrast, the single-CNN model generalized poorly: the matched fitting strategy performed substantially better in this case (A1: p = 4.93 × 10^−81^, PEG: p = 6.46 × 10^−33^).

Since it was fit to data from a single neuron, the single-CNN model was not likely to capture the sensory space encoded by a novel neuron. However, the held-out model did perform better than chance, so it provided a useful baseline to contrast with the ability of the population models to generalize. The finding that the pop-LN model generalized to new data illustrates a broader value of the population modeling approach. The LN model spans a more limited encoding subspace than the CNN models and thus does not predict activity as accurately. Still, this population model captures the auditory space spanned by the LN model, permitting the simpler architecture to generalize to data from new neurons. This observation suggests that the population modeling approach can benefit analysis using any model architecture, including new architectures that outperform those considered in the current study.

Given the ability of a pre-trained model to generalize to new data, we reasoned that such a model should also be beneficial to the analysis of smaller datasets that measure neural responses to fewer auditory stimuli. The amount of data available from neurophysiological recordings is often limited relative to the large datasets typically required for CNN models. This problem is especially acute in studies of animal behavior, where data is limited by the number of trials an animal is motivated to perform during a single recording session^33^. Thus, a pre-trained model that can accurately describe small datasets acquired in diverse experimental settings would be of broad value to the study of auditory coding.

To test for benefits of generalization on smaller datasets, we subsampled spiking data over 10-100% of the original dataset. A 1Dx2-CNN held-out model was pre-trained on 100% of data from all but one recording site, as above. The output layer was then re-fit to individual neuron responses from the excluded site, using only the subsampled data. In other words, the model drew on a much larger dataset for fitting the initial layers but only used the smaller subsample for fitting the final layer. We compared this model to a standard fit, in which both stages of fitting used data from all recording sites, but only a subsample of data was used for the entire fit. The standard model served as a control by representing a scenario in which data quantity is limited by experiment duration or other factors and no pre-trained model is available. Performance of models fit with the smaller datasets was quite variable, but the pre-trained held-out model performed better than the standard model on average (Fig. 6A, signed-rank test, p = 2.91 × 10^−21^). The benefit of pre-training extended across all subsamples tested (Fig. 6B, signed-rank test, p < 10^−9^).

**Figure 6.**
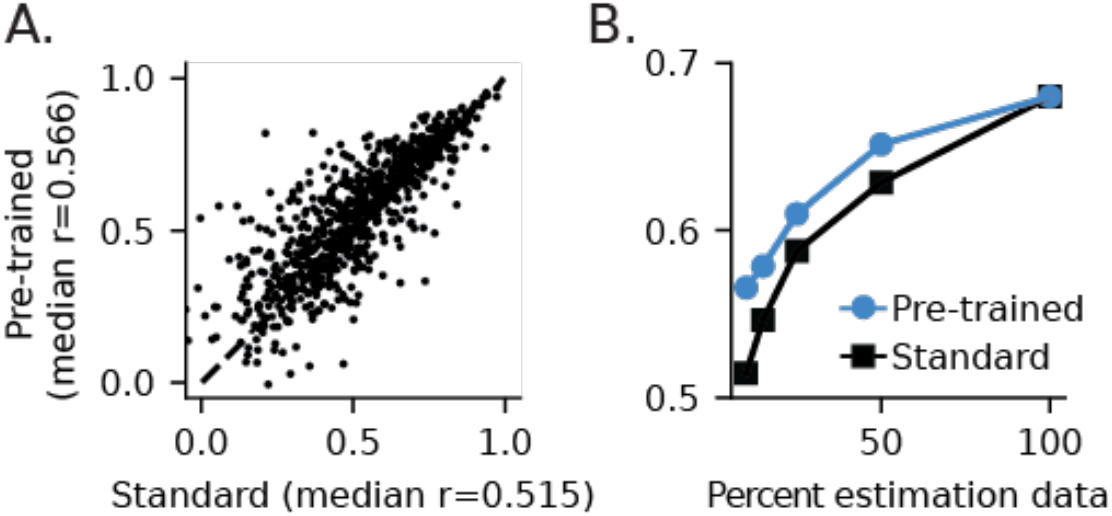
Generalization of a pre-trained 1Dx2-CNN model to smaller datasets. **A**. A pre-trained model was fit to every stimulus with neurons from one recording site excluded. The output layer was then re-fit for the excluded neurons, using a fraction of the available stimuli. The standard model was fit to all neurons, but only a subset of stimuli was used for the entire fit. Scatter plot compares prediction correlation between the pre-trained and standard models using 10% of available data. On average, the pre-trained model more accurately predicted the subsampled data (signed-rank test, p = 2.91 × 10^−21^). **B**. Curves compare median prediction correlation for pre-trained and standard models fit to subsampled data. Improved accuracy of the pre-trained model was consistent across all subsample sizes (signed-rank test, p < 10^−9^). Performance converges when 100% of data are used to fit both models.

### CNN models predict sparse cortical responses

A model that provides a truly general characterization of neural encoding should account for higher order properties of neural activity beyond basic selectivity for spectro-temporal features. One characteristic property of auditory cortex and other sensory brain areas is their high lifetime sparseness^25,34,35^. Lifetime sparseness measures the frequency at which stimuli evoke changes in spike rate from spontaneous levels. As sparseness increases, neurons become selective for progressively smaller subsets of stimuli. A sparse code has been proposed as an efficient strategy for exposing important features in complex natural stimuli^24^, but linear filters tend to predict responses with much lower sparsity than is actually observed^25^.

To evaluate the ability of CNN models to account for sparse activity in auditory cortex, we computed lifetime sparseness for responses predicted by the pop-LN and 1Dx2-CNN models, as well as for the actual response of the modeled neurons. Histograms of spike rate for an example neuron illustrate that the CNN model prediction matched the distribution of actual spiking more closely than the LN model (Fig. 7A-B). In this example, the CNN prediction histogram peaks at 0 spikes/sec, while the LN histogram peaks at 1 spike/sec. On average, the CNN model was less sparse than the actual data but predicted consistently sparser responses than the pop-LN model in both A1 and PEG (Fig. 7C, D; signed-rank test, A1: p = 5.30 × 10^−11^, PEG: p = 1.70 × 10^−12^). Analysis of the relationship between prediction correlation and predicted sparseness indicated that increased sparseness was driven by the improved performance of the CNN model (Supplementary Fig. 4). Thus, in addition to accounting for more variance in evoked activity, the CNN model was better able to capture a higher order property of neurons in auditory cortex.

**Figure 7.**
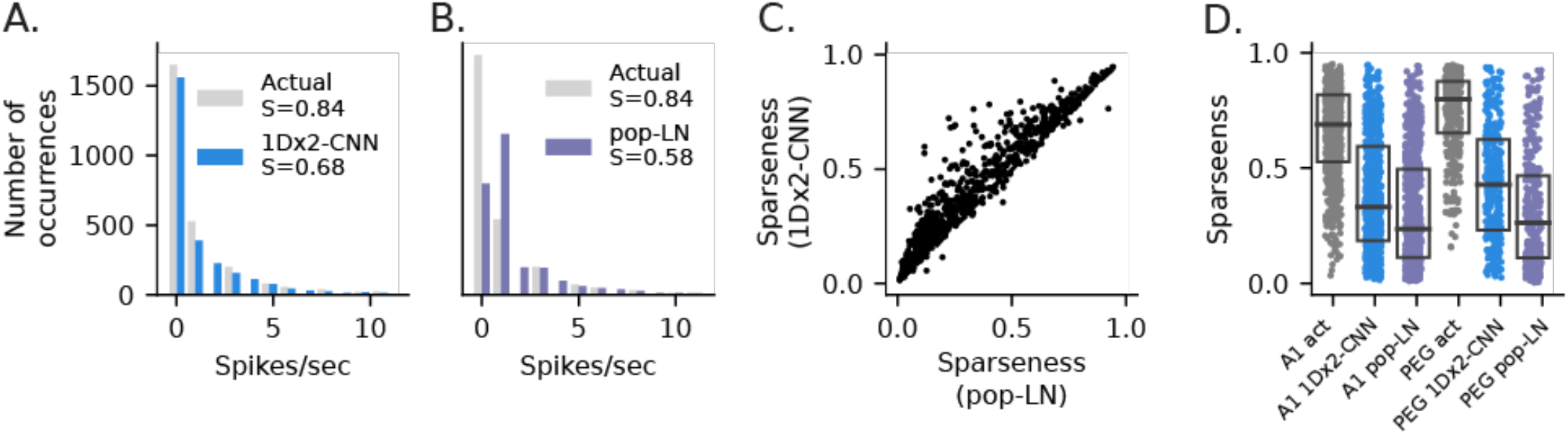
CNN models predict sparse auditory selectivity. **A**. Histogram of actual spike rate (gray) and rate predicted by the 1Dx2-CNN model (light blue) for one unit during the entire course of natural sound presentation in the validation data. Predicted and actual responses follow closely matched distributions, peaking at zero spikes/sec. **B**. Histogram of actual response and prediction by the pop-LN model, plotted as in A. The LN distribution peaks at 1 spike/sec and is less similar to the actual data. **C**. Scatter plot compares lifetime sparseness predicted by the pop-LN and 1Dx2-CNN models for auditory-responsive A1 neurons. CNN predictions were consistently sparser. **D**. Distribution of actual sparseness and sparseness predicted by the two models for A1 and PEG data. The 1Dx2-CNN model did not match the sparseness of the actual data but predicted greater sparseness than the pop-LN model (U test, A1: p = 5.30 × 10^−11^, n = 777; PEG: p = 1.70 × 10-12, n = 339).

## Discussion

We developed a convolutional neural network (CNN) architecture that simultaneously models the responses of several hundred neurons to dynamic natural stimuli. These population CNNs substantially outperformed traditional LN models of auditory coding, as well as CNNs fit to the activity of individual neurons. Moreover, population models were generalizable: the output layer of a pre-trained model could be re-fit to novel data without a reduction in prediction correlation. The generalizability and improved performance of the population CNN models is consistent with the hypothesis that their early layers describe a comprehensive spectro-temporal space encompassing sound encoding by any neuron in the auditory field being analyzed.

### Population network models provide a natural expansion of linear-nonlinear models

Alternatives to the LN model have been proposed to better characterize auditory encoding. Polynomial expansions, based on Taylor and Volterra series, are the classic extension of linear models to better account for sensory activity^12,36,37^. Although theoretically well motivated, these second-order models provide only modest improvements over the first-order, linear model. This shortcoming is likely due to two factors. First, second-order models require larger datasets for fitting and are thus more prone to estimation noise than linear models. Second, even with adequate fit data, the expanded functional space of a second-order model may not align well to the actual nonlinear biological properties of auditory neurons. Other alternatives attempt to address the latter limitation by accounting for specific biological nonlinearities, such as contrast gain control and short-term plasticity^3,14,38^. With an empirically targeted design, these models tend to require less data for fitting. However, they still only provide modest improvements to performance and may be limited in their ability to account for the full complement of nonlinearities present in sensory neural responses.

Artificial neural networks provide a third alternative that can account for a broader range of nonlinear properties. These large and complex models have traditionally required specialized fitting procedures to account for sampling limitations and noise in spiking data^15^. In the current study, we showed that the LN model architecture can be extended smoothly into small neural networks with convolutional units resembling linear filters in LN models. Moreover, we leveraged statistical power across neurons to fit these models on a modestly sized dataset without highly tuned optimization techniques. Population CNN models fit using this approach performed consistently better than the LN model and other nonlinear models^14^.

For all population CNN architectures tested (1D-CNN, 1Dx2-CNN, 2D-CNN), prediction correlation increased as model complexity increased and reached an asymptote at complexities of about 150-200 free parameters per neuron. This number reflects an increase in model complexity relative to an efficiently parameterized LN model, but still falls far short of the approximately 450 free parameters per neuron required for an equivalent full-rank LN model^31^. The relationship between fit dataset size and prediction correlation shows that performance was limited by the amount of fit data available. Thus, further increasing the amount of fit data should lead to even more accurate model predictions. A more general model that can account for this additional data will likely require more free parameters.

Surprisingly, there was not a clear “winner” among population CNNs: these models were highly equivalent, and differences in prediction correlation were small. Despite theoretical guarantees that *some* CNN exists that *can* approximate a given neural encoding function, there is no guarantee that any specific CNN of a fixed size can meet that goal. There is also no guarantee that a network’s parameters can be optimized for a particular function algorithmically, even if the chosen architecture is sufficiently large^39^. Accordingly, we had no expectation of equivalence between the CNNs we built despite their similarities. In practice, for example, one might expect idiosyncrasies of neural activity, such as non-Gaussian noise and a small fit dataset, to make some models more susceptible than others to fit errors or local minima. However, our results demonstrate that population CNN models can be fit robustly, regardless of details of the model architecture.

### Toward a generalized model of auditory cortex

The goal of a fully generalizable brain model that can simulate neural activity across novel experimental conditions is not new^40^. However, attempts to build detailed models from the ground up have met with limited success, and it remains unclear which details of neural circuits and biophysical mechanisms are required to mimic natural function. In the current study, we took a more functional approach to building a generalizable model. We used nonlinear regression to model the auditory system, without accounting for detailed biological circuitry. In this sense, the population CNN models we developed bear a closer resemblance to foundation models, which have grown increasingly valuable across the field of machine learning^20^. In the domain of language processing, for example, models such as BERT and GPT-3 can be applied to a wide range of language problems with little additional training^21,41^. A foundation model for biological auditory coding could draw on existing foundation models or the architectures developed here to describe a wide range of neurophysiological processes. Previous studies of auditory and visual neurophysiology have promoted a similar idea^16,19^. Intermediate representations in large CNN models, initially fit for a visual or image processing problem, can be used in a generalized linear model to predict neurophysiological responses to sensory stimuli. Here, we found such an approach was able to account not just for sensory selectivity but also for the time-varying response to dynamic natural stimuli.

A general model for biological auditory processing has wider applications than explaining selectivity for sound features. In this study, for example, a population CNN model also accounted for sparse coding properties in auditory cortex better than the LN model. This property emerged from the model even though it was not imposed by the fitting process. Thus, population models trained on larger and more diverse datasets should account for a wide range of functional properties, in the same way that foundation models generalize to applications outside the specific problem for which they were trained.

### Functional differences across the cortical hierarchy

One open question in studies of the auditory system is how sound representations evolve across the cortical processing hierarchy. Traditional LN models have been effective in the brainstem and midbrain^42–44^, but their accuracy is limited in auditory cortex^2,3^. Studies comparing multiple cortical fields are limited but indicate LN models perform even more poorly in non-primary auditory cortex than in A1^27,31^. This is expected since sound-evoked activity in non-primary cortex is generally less reliable, reflecting greater sensitivity to internal state^27,45^ and complex, long-lasting sensory adaptation that limits the efficacy of encoding model analysis^27,46^. In this study, CNN models were able to achieve a substantial improvement in performance over the LN model in a non-primary field (PEG). However, prediction correlation was consistently lower for PEG than for A1, even after accounting for differences in response reliability (SNR) between areas. This difference is consistent with the idea that PEG neurons exhibit more selective, nonlinear response properties than A1. In the absence of data limitations, a CNN or related model should be able to account for this sensory selectivity. Thus, the difference in performance indicates that a larger fit dataset is required to characterize PEG neurons as accurately as A1 neurons.

### Neural population dynamics and the space of sensory representation

Questions around the dimensionality of cortical sensory coding spaces are an active area of research^47^. The fact that population CNNs generalized readily to novel neural data in this study suggests that embedded in their parameters was a complete representation of the spectro-temporal features encoded across the entire cortical field. The dimensionality of a population model’s final hidden layer determines how many channels are recombined to predict the activity of any neuron. The best-performing 1Dx2-CNN model relied on 100 channels in this layer, and the performance of the held-out models indicates that the constrained space this layer represents could account equally well for the activity of any neuron in A1. This result suggests that much of the sensory activity of the many thousands of neurons in auditory cortex can be accounted for by a relatively low-dimensional space.

As high channel-count recordings in neurophysiological research continue to become more prevalent and grow in scale, the feasibility and value of population models will also increase. We expect these benefits to be particularly strong in circumstances where dataset size is constrained by experimental design, such as in studies of behavior. When behavioral factors such as motivation to perform a task limit the number of trials during recordings, statistical power can be increased by recording from a large number of neurons. In addition, sensory coding properties can be considered in a constrained sensory subspace by using a model pre-trained on a larger dataset.

## Methods

### Data collection

All procedures were approved by the Oregon Health and Science University Institutional Animal Care and Use Committee and conform to standards of the Association for Assessment and Accreditation of Laboratory Animal Care (AAALAC).

Prior to experiments, ferrets (*Mustela putorius furo*, n = 5) were implanted with a custom steel head post to allow for stable recording. While under anesthesia (ketamine followed by isoflurane) and under sterile conditions, the skin and muscles on the top of the head were retracted from the central 3 cm diameter of skull. Several stainless-steel bone screws (Synthes, 6 mm) were attached to the skull, the head post was glued on the mid-line (Charisma), and the site was covered with bone cement (Charisma and/or Zimmer Palacos). After surgery, the skin around the implant was allowed to heal. Analgesics and antibiotics were administered under veterinary supervision until recovery.

After animals recovered from surgery and were habituated to a head-fixed posture, a small craniotomy (approximately 0.5 mm diameter) was opened over A1 or the secondary auditory field, PEG, immediately ventro-anterior to A1^26,27^. Neurophysiological activity was recorded using silicon multielectrode arrays (UCLA probes^48^). The array was inserted approximately normal to the cortical surface using a microdrive (Alpha-Omega Engineering EPS). Electrophysiological activity was amplified and digitized (Intan RHD-128) and recorded using open-source data acquisition software (OpenEphys). Recording site locations were confirmed as being in A1 or PEG based on tonotopy, frequency tuning and response latency^26,27,49^.

Single- and multi-unit spiking events were extracted from the continuous, multichannel electrophysiological traces using Kilosort 2 ^50^. Units were only kept for analysis if they maintained isolation and a stable firing rate over the course of the experiment. Unit isolation was quantified as the percent overlap of the spike waveform distribution with neighboring units and baseline activity. Isolation > 95% was considered a single unit, and isolation > 85% was considered a multi-unit (single units: A1, 567/849; PEG, 314/398). There was no significant difference in median prediction correlation for any of the exemplar models between these groups in either brain area (U test, p > 0.05). Thus, we pooled single- and multi-unit data into a single population for this study, and we refer to these units as “neurons.”

Stimulus presentation was controlled by custom software written in Matlab (Mathworks, R2017A). Digital acoustic signals were transformed to analog (National Instruments PCI6259) and amplified (Crown D-75a). Stimuli were presented through a flat-gain, free-field speaker (Manger) 80 cm distant, 0-deg elevation and 30-deg azimuth contralateral to the neurophysiological recording site. Prior to experiments, sound level was calibrated to a standard reference (Brüel & Kjær). Stimuli were presented at 60–65 dB SPL (peak-to-peak amplitude).

### Natural sound stimuli

Data were collected during presentation of a library of natural sounds (595 1-sec samples, 0.5 sec ISI). Approximately 15% of these sounds were ferret vocalizations and environmental noises in the animal facility, recorded using a commercial digital recorder (44-KHz sampling, Tascam DR-400). Recordings included infant calls (1 week to 1 month of age), adult aggression calls, and adult play calls. No animals that produced the vocalizations in the stimulus library were used in the current study. The remaining 85% of sounds were drawn from a library of human speech, music and environmental noises developed to characterize natural sound statistics^51^. Activity was recorded during a single presentation of 577 samples and 20 repetitions of the remaining 18 samples. The low-repetition data were used for model estimation and the high-repetition data were used for model validation.

### Modeling framework

For all analyses used in this study, spike data for each neuron was converted into a peristimulus time histogram (PSTH), *r*(*t*), the time-varying spike rate, sampled at 100 Hz (10 ms bins). The input to each model consisted of a sound waveform converted into a spectrogram, *s*(*f,t*), using log compression and a second-order gammatone filter bank to account for the action of the cochlea^52,53^. This step was fixed for each model rather than fitting the spectrogram’s parameters, since we have observed little benefit from this additional complexity^31^. The filter bank included *F* = 18 filters with *f*_*j*_ spaced logarithmically from *f*_low_ = 200 to *f*_high_ = 20,000 Hz (approximately 1/3 octave per bin). The filter bank output was downsampled to 100 Hz to match the sampling of the neural PSTH. Filter shapes in the subsequent model definitions correspond to these 1/3-octave frequency bins and 10 ms time bins.

#### Linear-nonlinear models

The linear-nonlinear spectro-temporal receptive field (LN) model is widely used in studies of neural auditory coding^28,49,54^, and was used as a baseline for this study (Fig. 1A). The first stage of the LN model convolves a finite impulse response (FIR) filter, *h*, with the stimulus spectrogram to generate a linear firing rate prediction, *r*_lin_:

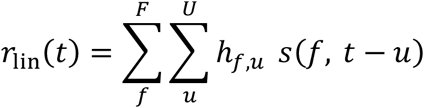

For the models used here, the filter consists of *F* = 18 spectral channels and *U* = 25 temporal bins. In principle, this transformation can be achieved with a single 18×25 filter. In practice, the filter was implemented as a rank D factorization: projection onto an 18xD spectral weighting matrix specified by a gaussian function followed by convolution with a Dx25 temporal filter, where D varied from 1 to 11. This implementation substantially reduces the number of free parameters without sacrificing model performance^31^.

A static sigmoid nonlinearity that mimics spike threshold and firing rate saturation is applied to the result of this convolution to produce the final model prediction. For this study, we used a double exponential nonlinearity:

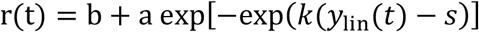

where the baseline spike rate, saturated firing rate, firing threshold, and gain are represented by *b, a, s* and *k*, respectively^31^. This model predicts the activity of each neuron independently from the rest of the recorded data, but we implemented it in TensorFlow using custom layers such that we could run a single “model” for the full neural population^55^. This approach enabled a dramatic speedup in fit time compared to a one-fit-at-a-time strategy and allowed us to use the same optimization routine as for the population models.

#### Single-neuron CNN models

Single-neuron CNN models (single-CNN, Fig. 1C) used in this study have two layers. The first layer is a 1D convolutional layer composed of many units that each apply the following operations in series: multiply Gaussian-distributed spectral weights with the spectrogram to produce a single channel, convolve this weighted channel with a rank 1, 250ms temporal filter, then apply an offset rectified linearity (ReLU) to the output of the convolution. The number of units in this layer determines the model’s size. We fit a total of six model variants in the current study, with unit count ranging from 2 to 18. The second layer consists of a single dense weighting unit with a double exponential nonlinearity as the activation function. The output of this unit is the model’s prediction of a single neuron’s time-varying firing rate.

The activation functions described here were also used for all other neural network models in this study: offset ReLU for all intermediate dense and convolutional layers unless otherwise specified, and double exponential (as described for the LN model) for the final dense layer (output).

#### Population CNN models

Population models are derived from the premise that neural populations cooperatively encode sensory information within a common subspace^7,56^. We implemented four population CNN architectures: a single 1D convolutional layer with no activation function followed by one dense layer (pop-LN, Fig. 1B), a single 1D convolutional layer followed by two dense layers (1D-CNN, Fig. 1D), two 1D convolutional layers followed by two dense layers (1Dx2-CNN, Fig. 1E), and three 2D convolutional layers followed by two dense layers (2D-CNN, Fig. 1F). For all population models, the number of units in the output layer is equal to the number of neurons in the population used for fitting (n = 849 for A1, n = 398 for PEG).

The pop-LN model resembles the LN model, in that the encoding properties of a single neuron can be collapsed into an LN model. The only difference is that the entire population shares a subspace defined by the convolutional layer, while weights for individual neurons are only computed in the final layer. We consider this design to be the minimal change needed to convert the single-neuron LN model into a population model. We tested 15 model variants within this architecture, where the number of convolutional units for each variant ranged from 4 to 300. Note that while the pop-LN model is technically a CNN, we refer to it as the LN model when contrasting it with the CNN models that contain intermediate nonlinearities.

The 1D-CNN model is similar to the single-CNN model, except there is a hidden dense layer after the convolutional layer. We compared sixteen model variants for this architecture, with 5 to 230 convolutional units and 10 to 300 hidden units.

The 1Dx2-CNN model is akin to the 1D-CNN model but has two consecutive convolutional layers instead of one. The first layer uses 150ms filters while the second uses 100ms filters, yielding the same 250ms total “context memory” as the models described above. We fit a total of 14 model variants for this architecture. The number of units in the first and second layers varied from 5 to 150 and 10 to 200, respectively, and the number of units in the hidden dense layer ranged from 20 to 250.

The 2D-CNN model also resembles the 1D-CNN model, but the 1D convolutional layer is replaced with three consecutive 2D convolutional layers each using ten 3×8 filters, encompassing a cumulative 240ms of context memory. We compared 15 model variants for the 2D-CNN architecture, where the number of units in the hidden dense layer ranged from 4 to 300.

To choose the structure of each model, we built progressively larger models out from the canonical single-neuron LN model by hand-selecting models with increasing size and number of network layers. For each model type, we evaluated a small number of wide-ranging hyperparameter combinations like learning rate and early stopping criteria, selected a combination that provided stable fits and good performance for all architectures, and then varied convolutional filter count and/or dense unit count to produce a continuum of model sizes. This was not an exhaustive exploration of the possible hyperparameters for the model architectures chosen, nor did we dive deeply into the full range of possible architectures. Additionally, we chose not to explicitly imitate biological nonlinearities like short-term plasticity or gain control within the models since the relatively brief stimuli used for this study were not designed to probe these slower response dynamics.

### Model optimization

Model parameters were fit using TensorFlow’s implementation of the Adam algorithm for stochastic gradient descent, using a mean squared error (MSE) loss function^30,55^. Post-fitting performance was measured on a separate validation dataset using prediction correlation, defined as the correlation coefficient (Pearson’s *R*) between model prediction and actual response, adjusted for uncertainty due to the finite sampling of validation data^32^.

For each fit, model parameters were randomly initialized nine times, with a tenth initialization set to the mean of the parameter distributions. Each initialization was used to fit a subset of the model with the output nonlinearity excluded. The best submodel fit was then used to initialize a subsequent full model fit. In past studies we found that this heuristic approach improved single-neuron model performance, and the same proved to be true for this study’s population models^14,31^. To mitigate overfitting, an early stopping criterion was set using twenty percent of the estimation data. We also tested dropout and L2 regularization^57,58^, but we found little to no benefit and chose to exclude them from this study for simplicity.

This fit procedure was consisted of two stages:

- First stage: Models were fit to all units from one brain region simultaneously (n = 849 A1; n = 398 PEG).
- Second stage: The output layer of each model was re-fit to one unit at a time while parameters for earlier layers were kept fixed.

The second stage served two purposes. First, it provided a modest increase in performance for population models by optimizing a separate loss function for each unit, rather than a single loss function produced by a weighted sum of the loss across neurons. Second, it ensured there were no differences in fitting process for the models in the generalization analyses.

For the test of generalization, the same two-stage procedure was used but with subsets of the data excluded. In held-out models, all K neurons from one site, S, were excluded from the first stage fit. Model parameters from this fit were used to initialize second stage fits for the K excluded neurons. In the case of the matched model, K neurons from other sites with prediction scores similar to those in site S (for a single-neuron LN model) were excluded during the first stage. Thus, the matched model provided a control for the generalization test, in that the number of neurons used for fitting this model was the same as for the held-out model. This method was repeated for every recording site to generate held-out and matched model predictions for every neuron.

### Exemplar models

After our exploration of a wide range of model sizes (Fig. 3A, B), subsequent analyses focused on exemplar models from each architecture. These exemplars were chosen such that each had similar complexity, and that complexity was as low as possible while keeping prediction correlation near the observed asymptote. Architectural hyperparameters were as follows, in layer-order, where N represents the number of neural responses in the data:

- LN: 1 1D convolutional unit (rank-5 factorization), 1 output unit.
- pop-LN: 120 1D convolutional units (250 ms), N output units.
- single-CNN: 6 1D convolutional units, 1 output unit.
- 1D-CNN: 100 1D convolutional units (250 ms), 120 hidden dense units, N output units.
- 1Dx2-CNN: 70 1D convolutional units (150 ms), 80 1D convolutional units (100 ms), 100 hidden dense units, N output units.
- 2D-CNN: 3 layers of 10 2D convolutional units each, in series (each 80 ms x 3 spectral bins, approximately 1 octave), 90 hidden dense units, N output units.

### Analysis of lifetime sparseness

Previous studies have argued that the brain encodes sensory stimuli with a sparse code, which provides a useful scheme for reading out sensory information downstream^24,25,34,35^. We used a lifetime sparseness metric to determine how well models were able to predict the sparsity of single units. The high repetition-count validation data provided a measure of sound evoked activity at each point in time, *r*(*t*), and sparseness was computed from the distribution of these responses^34^:

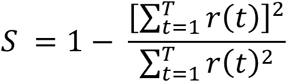

According to this metric, neurons with zero response at most times and a non-zero response for only a few times will have high sparseness (*S* = 1), while neurons with many non-zero responses will have low sparseness (S = 0). To evaluate predicted sparseness, time-varying model predictions were subjected to the same metric.

### Statistical methods

For all statistical comparisons, we used non-parametric tests since the distributions of the relevant variables were non-normal and no other distribution was apparent. For paired tests, e.g., comparing prediction correlations between different models on the same set of neurons, we used a two-sided Wilcoxon signed-rank test (referred to as “signed-rank test”). For non-paired tests, we used a two-sided Mann-Whitney U test (“U test”). Statistical significance was assigned for p < 0.05. Full p-values are reported for completeness unless many tests are reported simultaneously. Note that due to the large number of units in the recordings, the p-values reported here are often extraordinarily small. These values reflect the relatively large number of neurons in the dataset, which provided substantial statistical power.

To determine whether a model’s prediction was above chance for a given neuron, or if one model’s prediction was significantly more accurate than another’s for that neuron, we used a jack-knifed t-test. For the population analyses reported in Figs. 3-10, we only included data from neurons for which the 1Dx2-CNN, pop-LN, and single-CNN exemplar models all performed above chance. We considered neurons that met this criterion to be auditory-responsive (n = 777/849 A1 neurons, n = 339/398 PEG neurons) and assumed the excluded neurons were non-auditory.

### A toolbox for systematic comparison of encoding models

All models in this study were fit using the Neural Encoding Model System (NEMS, https://github.com/LBHB/NEMS). We developed this open-source software package to be a flexible and extensible tool for fitting models to sensory neurophysiology data. Implementing all models in a common framework helped eliminate potentially problematic differences in our analysis pipeline, like optimization routine and cost function evaluation, that otherwise might arise. We will continue expanding the model architectures supported by NEMS, and we invite contributions and suggestions to make this software helpful to the broader neuroscience community.

## Acknowledgments

The authors would like to thank Charles Heller and Mateo Lopez Espejo for assistance with data collection. We would also like to thank Nima Mesgarani and Menoua Keshishian for their advice on fitting CNN models.

## Author contributions

J.P. performed the majority of data analysis and software development. Both J.P. and S.D. designed the study and wrote and edited the manuscript.

