## Supplemental Materials for "Can deep learning provide a generalizable model for dynamic sound encoding in auditory cortex?"

JRP ORCID iD: https://orcid.org/0000-0001-9393-0474

SVD ORCID iD: https://orcid.org/0000-0003-4135-3104

### Contents

(Pg 2): Supplementary Figure 1: Predicted and actual PSTHs for two example neurons.

(Pg 3): Supplementary Figure 2: Model equivalence.

(Pg 4): Supplementary Figure 3: SNR control and cross-region generalization.

(Pg 5): Supplementary Figure 4: Lifetime sparseness as a function of model performance.

(Pg 6): Supplementary Methods: Model equivalence.

(Pg 7): Supplementary Methods: Reliability of neural responses (SNR).

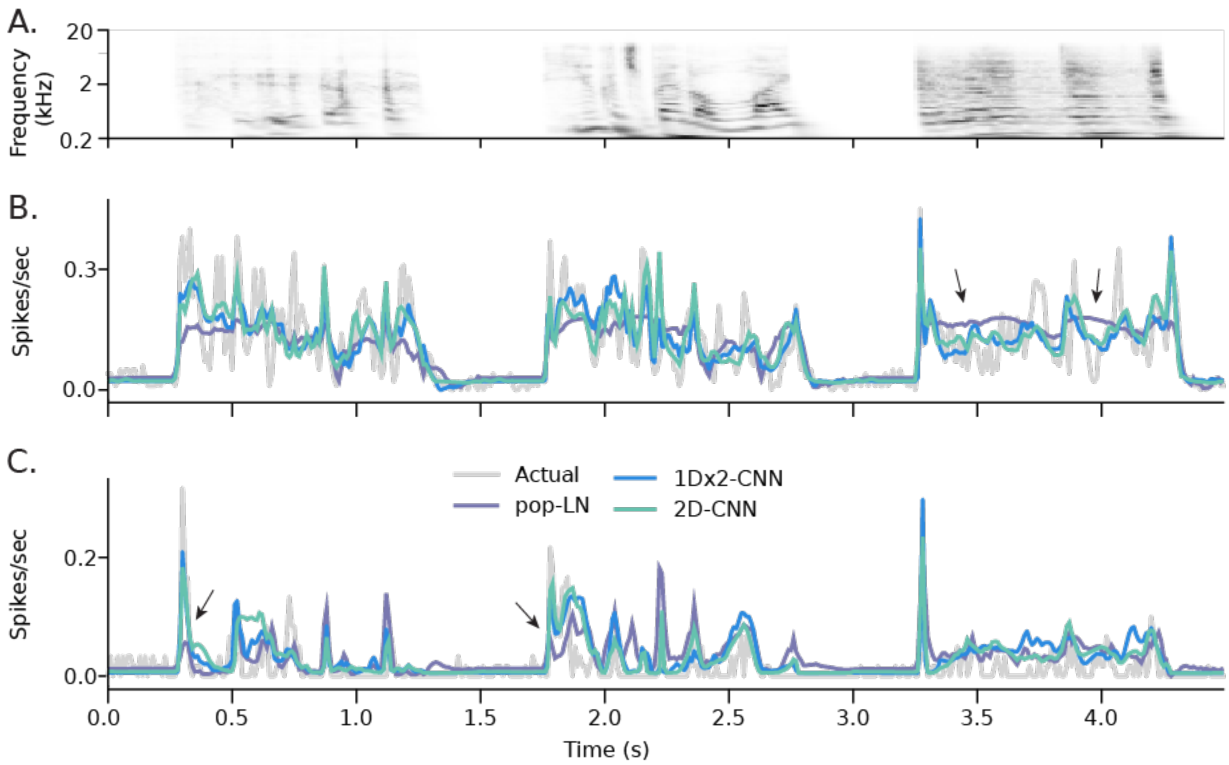


**Supplementary Figure 1.** Examples of actual versus predicted PSTHs for two A1 neurons. **A.** Segment of the stimulus spectrogram from the validation dataset used for testing prediction accuracy. **B.** Actual PSTH (gray) overlaid with the PSTHs predicted by the 1Dx2-CNN (blue), 2D CNN (light green), and pop-LN (purple) models. The two CNN models produced similar predictions that correlated well with the actual response (1Dx2-CNN: r = 0.812, 2D CNN: r = 0.800). The pop-LN model failed to predict many of the temporal features in the actual PSTH (r = 0.590, arrows at 3.5, 4.0 sec). C. Actual and predicted PSTH for a second neuron, plotted as in B. Again, both CNN models had higher prediction correlation. Here their accuracy is evident in the observation that they accounted for large transient responses better than the LN model (1Dx2-CNN: r = 0.729, 2D CNN: r = 0.748, pop-LN: r = 0.535; arrows at 0.25, 1.75 sec).

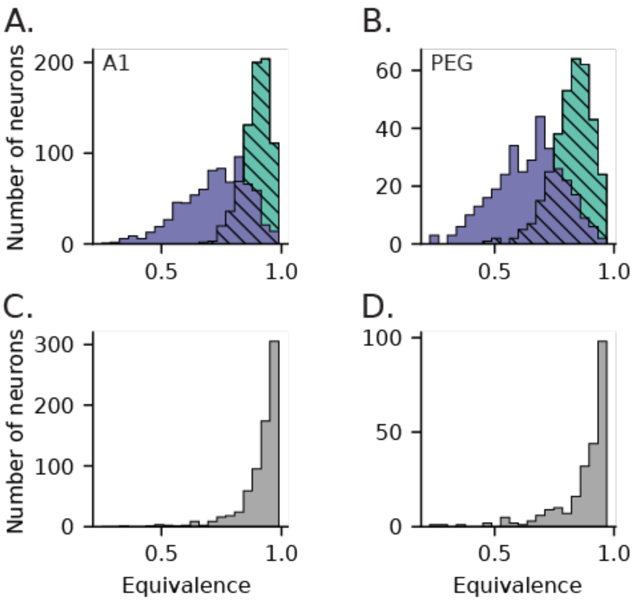


**Supplementary Figure 2.** Quantification of equivalence between CNN, pop-LN, and LN exemplar models. **A**. Histogram of equivalence (correlation between predicted PSTHs, see Supplementary Methods) on the validation data for auditory-responsive A1 neurons (n = 777/849), between 2D CNN and 1Dx2-CNN models (light green, hatched) and between 1Dx2-CNN and pop-LN models (purple). Equivalence was greater between the two CNN models than between the 1Dx2-CNN and pop-LN models (signed-rank test, $p = 1.47 \times 10^{-128}$). This result indicates that CNN models achieved higher prediction accuracy over the LN architectures in similar ways. **B.** As A, but for PEG neurons (n = 339/398). Here again we observed higher median equivalence between the CNN models ($p = 1.64 \times 10^{-55}$). **C.** Histogram of equivalence between LN and pop-LN models for A1 neurons. The distribution is shifted even farther toward the 1.0 bound, indicating that the LN and pop-LN models predicted closely matched PSTHs for most neurons. **D.** As C, but for PEG neurons. The equivalence distribution is similarly right shifted.

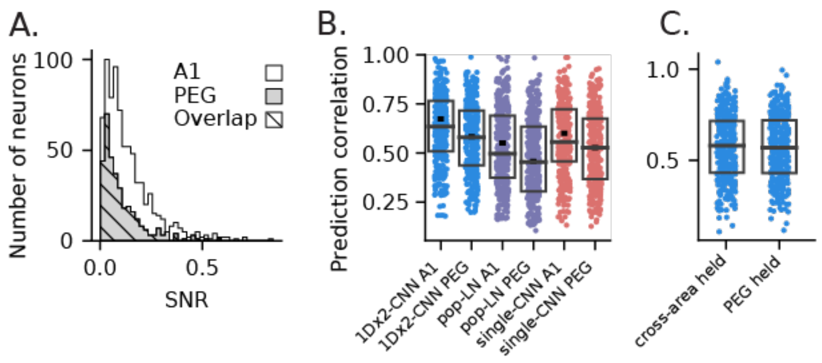


**Supplementary Figure 3**. Comparison of model performance between primary (A1) and secondary (PEG) auditory cortical regions after controlling for differences in SNR. **A**. SNR scores (see Supplementary Methods) for all neurons from A1 (white), PEG (gray), and an overlapping subset (hatched) of neurons with matched distributions. Median SNR was higher for A1 (md = 0.0989, U test: p = 2.49 x 10-6) than for PEG (md = 0.0713), and the A1 subset had a lower median SNR (md = 0.0709) while the median for the PEG subset was mostly unchanged (md = 0.0710). To form the overlapping set, we selected the largest possible subset of neurons from each brain region for which binned distributions of SNR scores for the two subsets were identical. With this approach, the subset was formed primarily by excluding high-SNR neurons from A1 since PEG neural responses were less reliable overall. **B**. Prediction accuracy of three exemplar models for neurons in the A1 and PEG subsets. Boxes show the 1st, 2nd and 3rd quartile for the selected subset, and the small horizontal dash indicates median performance across all neurons from that brain region. The increase in prediction accuracy for A1 was smaller for this subset, but still significant (U test; 1Dx2-CNN: p = 6.685 x $10^{4}$; pop-LN: p = 6.743 x $10^{-4}$; single-CNN: p = 4.78 x $10^{-4}$), indicating that the increased SNR of A1 responses only partially accounts for our models' higher prediction accuracy for A1 neurons. **C.** Prediction correlations for the 1Dx2-CNN PEG held-out model (Fig. 5C, median r = 0.569) and a "cross-area" held-out model (md = 0.581). To fit the cross-area model, we used the held-out approach (Fig. 5A) to pre-train a 1Dx2-CNN model on A1 data, and then fit the output layer using data from PEG neurons in the matched-SNR subset. The important distinction in this case is that the output layer of each model was always fit to data from a PEG neuron, but the earlier layers were fit using an independent set of either A1 or PEG responses. Pre-fitting to A1 data proved to be just as effective as pre-training on PEG data for predicting PEG neural responses (signed-rank test, p = 0.119). This similarity in performance suggests that CNN models generalized across cortical areas, in addition to new neurons in the same area.

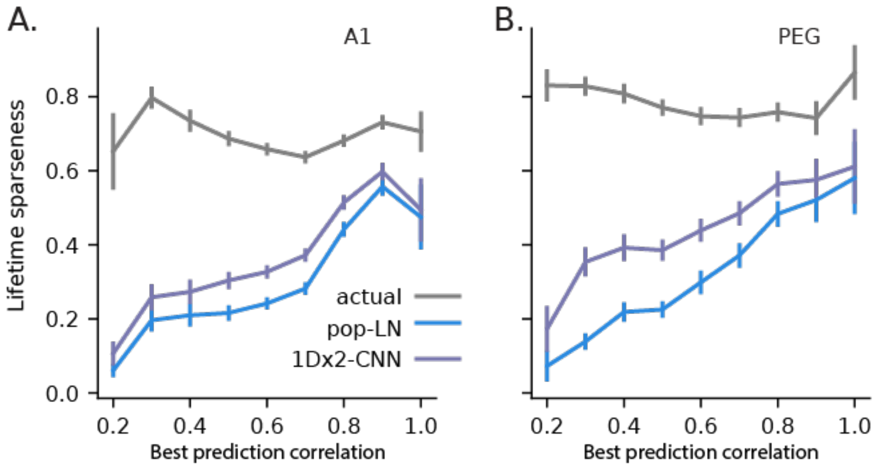


**Supplementary Figure 4.** Mean actual and predicted sparseness as a function of model prediction accuracy. **A.** Lifetime sparseness was computed for the time-varying neural response and the response predicted by pop-LN and 1Dx2-CNN models for each neuron in the A1 validation dataset. Average sparseness values were computed after binning by the best prediction correlation for those two models. Sparseness of predicted activity consistently increased with prediction correlation. Error bars indicate 1 SEM. **B.** Comparison of actual and predicted sparseness for PEG, plotted as in A.

**Supplementary Methods: Model equivalence**

A striking observation from our performance analysis is that the 1Dx2-CNN and 2D CNN models predicted activity with nearly equal accuracy (Fig. 4B). Given the differences in architecture, it was not immediately clear whether both models captured the same functional properties or if their improvements over the LN model instead reflected their ability to capture distinct aspects of the neurons' function. Qualitative inspection of PSTHs predicted by the 1Dx2-CNN and 2D CNN models suggests that they produced similar predictions, which were visibly distinct from the pop-LN model prediction (Supplementary Fig. 1).

To quantify prediction similarity between models, we computed model "equivalence" on the validation data as the Pearson correlation coefficient between PSTHs predicted for each neuron. This approach enables a relative comparison: if one distribution of equivalence scores is shifted farther right, then that pair of models is more similar than the pair of models with the leftmost distribution. This analysis was a variant of a previous method that computed the partial correlation between PSTHs, relative to the prediction of a baseline model[1]. The simpler comparison used in this study introduces the limitation that it cannot be meaningfully applied to models that explain a high amount of variance in the data. In that case, the model predictions *must* be highly correlated with each other, so a separate baseline model would be required to measure their similarity.

**Supplementary Methods: Reliability of neural responses (SNR)**

Sensory responses of PEG neurons tend to be noisier, or less reliable, than A1 responses[2], which could explain the observed difference in prediction accuracy between the two brain regions (Figs. 3, 4). To test this possibility, we generated subsets of data from A1 and PEG with balanced levels of auditory responsiveness by assessing the reliability of auditory responses across repeated presentations of the same stimulus, using an approach modified from a previous study[1]. We refer to this reliability quantification as SNR and define it as follows. We define *total power*, *T*, as the variance of the time-varying spike rate, $r_j$, averaged across presentations of the same stimulus, $j = 1 \ldots m$, for a given neuron,

$$\mathrm{T} = \frac{1}{m}\sum_{k=1}^{m}\langle r_j, r_j\rangle$$

where $\langle \ldots \rangle$ denotes a dot product. We define *signal power*, *S*, as the average correlation between the time varying spike rate on different presentations of the same stimulus:

$$\mathrm{S} = \frac{1}{m}\sum_{k=1}^{m}\langle r_j, r_k\rangle,\ j \neq k$$

This process is repeated for each stimulus $i = 1 \ldots n$. SNR is then measured as the average ratio of signal power to total power across stimuli,

$$\mathrm{SNR} = \frac{1}{\mathrm{n}}\sum_{\mathrm{i}=1}^{\mathrm{n}}\frac{S_i}{T_i}$$

### Supplementary Materials References


1. Pennington, J. R. & David, S. V. Complementary Effects of Adaptation and Gain Control on Sound Encoding in Primary Auditory Cortex. *eNeuro* **7**, (2020).
2. Atiani, S. *et al.* Emergent selectivity for task-relevant stimuli in higher-order auditory cortex. *Neuron* **82**, 486–499 (2014).
